# Investigating temperature effects on coastal microbial populations and trophic interactions with 16S and 18S rRNA metabarcoding

**DOI:** 10.1101/2021.03.17.435717

**Authors:** Sean R. Anderson, Margot Chisholm, Elizabeth L. Harvey

## Abstract

Temperature is a universal driver of microbial life, with rising sea surface temperatures expected to differentially influence the physiology, biodiversity, and distribution of bacteria and plankton. The impact of ocean warming on microbial interactions remains unclear, despite the importance of these relationships for ecosystem functioning. We employed weekly to monthly 18S and 16S rRNA gene amplicon metabarcoding over a full year (33 d) in a subtropical estuary, investigating microbial population dynamics and network interactions with respect to a temperature gradient (9–31°C). Certain microbes (e.g., Acidimicrobiia, Nitrososphaeria, and Syndiniales) increased in relative abundance with rising temperatures (Spearman ρ > 0.69), whereas other groups (e.g., Alpha- and Gammaproteobacteria, Bacillariophyta, and Dinophyceae) slightly decreased, became saturated, or remained stable. With network analysis, we observed an increase in 18S– 18S interactions in warm (23–31°C) vs. cold (<23°C) temperatures, largely involving Syndiniales, Bacillariophyta, and Dinophyceae ASVs. Bacteria ASVs were more connected to other microbes (higher degree and centrality) and became more prominent in the cold network, highlighted by well-established cross-domain relationships (e.g., diatom–bacteria) and positive interactions among bacteria (e.g., SAR11 and Rhodobacterales). These efforts highlight the types of interactions that may be more common under changing temperatures, with implications for modeling biogeochemistry and assessing ecosystem health.

## Introduction

Microbes (e.g., bacteria, archaea, and protists) form the base of marine food webs and are responsible for many important ecological and biogeochemical processes, such as carbon fixation, trophic transfer, nutrient cycling, and organic matter remineralization (Azam *et al.*, 1983; Caron and Hu, 2019). Microbial communities are not static, instead characterized by diverse and complex interactions at the cellular level (e.g., predator–prey or symbiont–host) that structure population-level dynamics and underpin ecosystem functioning (Worden *et al.*, 2015). Microbes and their trophic interactions are sensitive to shifts in the surrounding physical and chemical environment (Boyd and Hutchins, 2012; Caron and Hutchins, 2013), with many of these environmental conditions rapidly changing due to human-derived CO2 emissions (Hutchins and Fu, 2017). Over the past century, atmospheric CO2 concentrations have increased by ∼50% (currently >400 ppm), leading to increased absorption of CO2 at the ocean’s surface and altering seawater properties like pH, temperature, nutrients, and oxygen (Doney *et al.*, 2020). Assessing the impacts of environmental change on microbial population dynamics and trophic activities remains critical, as this information will guide predictions on the response of microbes to future ocean conditions, as well as the effects on carbon transfer, biogeochemistry, and ecosystem resources (e.g., tourism and fisheries).

Rising surface water temperatures represent one of the most well-documented consequences of global climate change, with temperatures expected to rise by ∼4°C over the next century (Pachauri *et al.*, 2014). Temperature is a universal driver of microbial growth and physiology (Eppley, 1972; Toseland *et al.*, 2013), and so, continued efforts have been made to characterize effects of ocean warming on microbes (Cavicchioli *et al.*, 2019). Among phytoplankton, measurements of temperature-based growth rates have revealed species- to strain-level differences in thermal response curves and acclimation (Barton and Yvon-Durocher, 2019; Anderson and Rynearson, 2020; Strock and Menden-Deuer, 2020), with implications for phytoplankton diversity, distribution, and bloom dynamics (Thomas *et al.*, 2012; Boyd *et al.*, 2013; Ibarbalz *et al.*, 2019). Some studies predict that ocean warming and associated effects (e.g., reduced mixing or nutrients) may favor smaller phytoplankton, like cyanobacteria or picoeukaryotes (Hare *et al.*, 2007; Feng *et al.*, 2009), over larger diatoms (but see Kling *et al.*, 2020) or promote formation of harmful blooms (Paerl and Huisman, 2009), all likely to influence ecosystem-wide processes.

Despite these insights, distinguishing the impacts of ocean warming on microbial life has been challenging, as microbes are functionally diverse and form complex trophic interactions that change over space and time (Ward *et al.*, 2017). For example, while higher temperatures are thought to directly impact heterotrophic bacteria production and respiration (Hutchins and Fu, 2017), there may also be indirect effects on bacterial populations, driven by temperature-related shifts in the availability and sources of phytoplankton-derived organic matter (Thornton, 2014). Warming may induce stronger phytoplankton-bacteria coupling (von Scheibner *et al.*, 2014), as evidenced by increased attachment and nutrient exchange between groups, suggesting a preference for mutualistic relationships under higher temperatures (Arandia-Gorostidi *et al.*, 2017a). More recent work observed an increase (up to 6-fold higher) in carbon assimilation under elevated temperatures for two copiotrophic bacteria (Flavobacteria and Rhodobacteraceae) associated with coastal diatoms, further indicating potential temperature-related changes to phytoplankton–bacteria interactions (Arandia-Gorostidi *et al.*, 2020). Other types of microbial interactions, such as predation of phytoplankton by heterotrophic protists, may also increase under warmer conditions (Chen *et al.*, 2012), though like primary producers, grazers exhibit species-specific variability in thermal acclimation and temperature-based growth strategies (Franzè and Menden-Deuer, 2020). Therefore, in addition to physiological impacts, increased temperatures are also likely to influence how microbes interact with each other, which will require further insight into how such interactions vary over natural temperature gradients.

Coastal marine regions, like estuaries, represent ideal locations to explore temperature-related shifts in microbial community dynamics and interactions, as temperature (and other abiotic factors) vary over hourly to monthly time scales and drive episodic phytoplankton blooms (Cloern *et al.*, 2014; Needham and Fuhrman, 2016; Needham *et al.*, 2018). Amplicon metabarcoding coupled with high-throughput sequencing has become increasingly used to monitor microbial diversity and composition across marine systems (Caron and Hu, 2019), and in estuaries, these techniques often identify greater species richness compared to microscopy or pigment analysis (Abad *et al.*, 2016; Gong *et al.*, 2020). Applying metabarcoding tools over space and time in these dynamic regions has enabled researchers to better explore how microbial communities respond to environmental fluctuations.

To establish temperature-based changes in microbial population dynamics and interactions, we employed weekly to monthly sampling of surface water over a full year in the Skidaway River Estuary (GA, USA), simultaneously targeting the response of major bacteria, archaea, and protist groups with 18S and 16S rRNA metabarcoding. In our previous work on this time series, the 18S data were analyzed, revealing temporal trends in protist parasites and clustering of protist communities at 23–31°C (Anderson and Harvey, 2020). In this study, we analyze the 16S data and include previously analyzed 18S data to establish group-specific changes in relative abundance with respect to a wide temperature gradient (9–31°C) and between distinct temperature conditions (<23 vs. 23–31°C), as identified in Anderson and Harvey (2020). Metazoans were also targeted with 18S primers and included here (additional 743 ASVs) to explore the impact of temperature on higher trophic levels, including taxonomic groups (e.g., copepods) known to consume protists (Steinberg and Landry, 2017). Using 18S and 16S sequences, we generated temperature-based covariance networks that were inclusive of cross-domain interactions, revealing differences in network structure and microbial interactions at temperatures <23 vs. 23–31°C. Incorporating ‘omics surveys into current biological monitoring will facilitate more holistic and accurate predictions on the fate of microbes to ocean warming and downstream impacts on ecosystem functions and resources.

## Results

### Environmental conditions and kingdom level diversity

Surface temperatures over the year ranged from 9–31°C (Fig. 1A). Total dissolved nutrients ranged from 19.09–141.44 μM and strongly covaried with temperature (Spearman ρ = 0.86), while bulk chlorophyll concentration exhibited a narrow range (1.32–6.39 μg L^-1^; Fig. 1A). Eukaryotic community composition (via NMDS) was structured based on two distinct temperature conditions (ANOSIM R = 0.74, *p* = 0.001; Fig. 1B), referred to here as cold (<23°C) vs. warm (23–31°C) conditions. For 18S, average observed richness (number of ASVs) and Shannon diversity were not significantly different between cold and warm conditions (*p* > 0.2; Fig. 1C). Unlike eukaryotes, 16S composition did not exhibit structure based on temperature, with the NMDS ordination failing to converge (stress = 0.26). Yet, mean 16S richness and Shannon diversity were significantly higher (*p* < 0.001) in warm compared to cold conditions (Fig. 1D). Species richness vs. sequence abundance curves were saturated across 16S and 18S, indicating an appropriate sequencing depth was reached (Fig. S1).

**Fig. 1:**
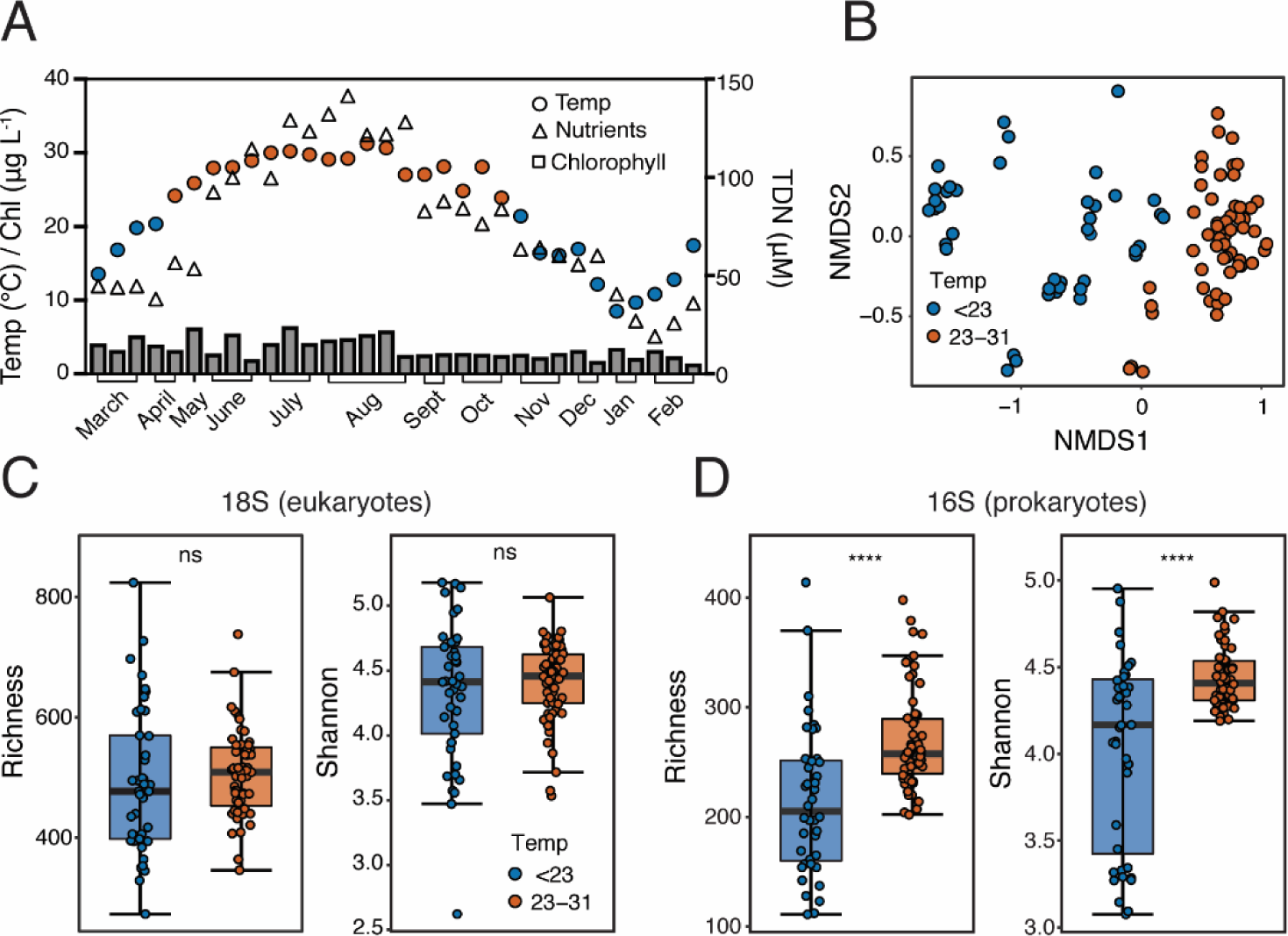
Key environmental parameters and temperature-based diversity estimates. **A**. Environmental conditions over the year in the estuary (adapted from Anderson and Harvey, 2019), including temperature (°C), chlorophyll concentration (μg L^-1^), and total dissolved nutrients (TDN = SiO_4_ + NO_3_ + PO_4_; μM). Samples were distinguished by temperature: cold (blue; <23°C, n = 42) vs. warm (red; 23–31°C, n = 57). **B**. Non-metric dimensional scaling (NMDS) of 18S communities under cold and warm conditions (stress = 0.12). NMDS for 16S failed to converge (stress = 0.26). **C, D.** Boxplots displaying mean number of ASVs (observed richness) and Shannon alpha diversity for 18S (**C**) or 16S (**D**) between cold and warm samples. Statistical differences between temperature are shown (ns = not significant, **** = p-value < 0.0001). Sample number is conversed across diversity plots.

### Temperature effects on group-specific abundance

To observe group-specific temperature effects on microbial abundance, we focused on 16S and 18S groups with the highest relative abundance on average (defined here as ≥1%) across all sampling points (Fig. 2). For 16S, the most abundant bacteria groups (on average) at the class level included Alphaproteobacteria (38%), Gammaproteobacteria (21%), Bacteroidia (15%), and Acidimicrobiia (10%), as well as less-abundant archaea groups like Thermoplasmata and Nitrososphaeria (each 1%; Fig. 2). We observed group-specific differences in 16S relative abundance vs. temperature relationships (Fig. 2). For instance, significant and non-linear relationships (*p* < 0.05) were observed for Alphaproteobacteria, Gammaproteobacteria, Bacteroidia, and Actinobacteria, with relative abundances either saturating and/or decreasing with rising temperatures (ρ = –0.24 to –0.76). Other groups like Acidimicrobiia and Deltaproteobacteria exhibited a more linear increase in relative abundance with temperature (ρ = 0.81 and 0.69, *p* < 0.001), while abundances of major archaea and cyanobacteria groups (e.g., Nitrososphaeria and Oxyphotobacteria) increased non-linearly with temperature (ρ = 0.8 and 0.82, *p* < 0.001; Fig. 2). Unlike other groups, relative abundance of Verrucomicrobiae was not significantly correlated with temperature (*p* = 0.32; Fig. 2).

**Fig. 2:**
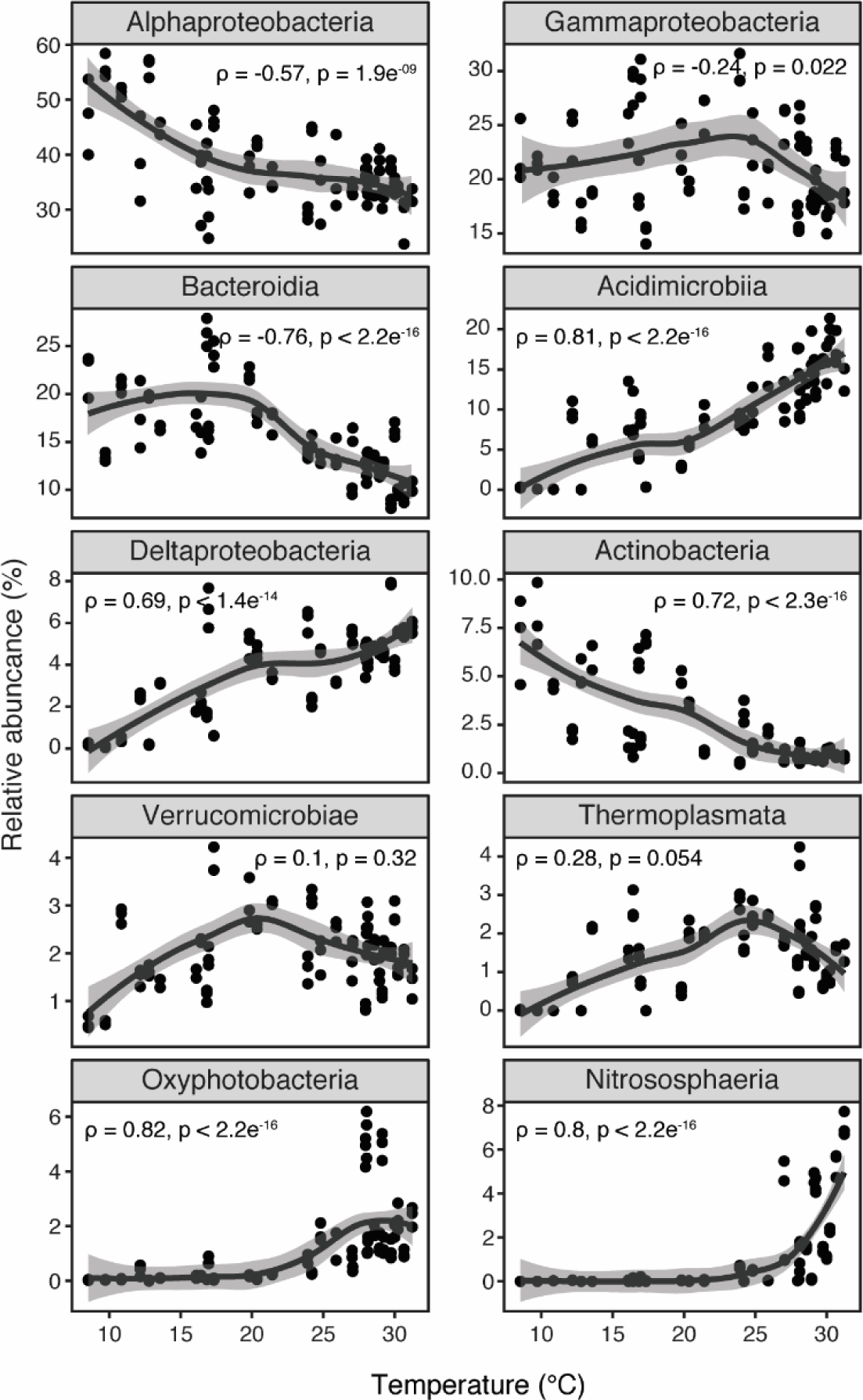
Group-specific temperature effects on 16S relative abundance. Relationships between 16S relative abundance for major groups (faceted by class) vs. temperature across all points. Local regression (loess) curves were fit to the data, with solid black lines representing smoothed trends and shaded gray regions the 95% confidence intervals of the fit. Spearman correlation coefficients (ρ) and *p*-values are indicated in the panels. Only the most abundant 16S (or 18S) class level groups (present in ≥1% of samples on average) were included (facets ordered by highest-lowest abundance). Triplicate samples shown for each temperature value.

Only 76 16S ASVs from the full dataset (0.05% of total) had significantly different (*p* < 0.001) log2 fold abundance between cold and warm conditions, with 69 of these ASVs belonging to major groups (Table S2). Log2 fold abundances exhibited a narrow gradient across 16S ASVs (often >20 or <–20), though group-specific differences were observed (Fig. S2). For instance, order level groups like SAR11 Clade and Rhodobacterales (Alphaproteobacteria), SAR86 Clade (Gammaproteobacteria), and Flavobacteriales (Bacteroidia) had differentially abundant ASVs under both warm and cold conditions (Fig. S2). In contrast, ASVs from Actinomarinales (Acidimicrobiia), Synechococcales (Oxyphotobacteria), and Nitrosopumiales (Nitrososphaeria) were observed only under warmer conditions (Fig. S2).

For 18S, the most abundant class level groups on average included protist assemblages such as Bacillariophyta (19%), Dinophyceae (17%), Mamiellophyceae (15%), and Syndiniales (9%), as well as several metazoan groups like Arthropoda (9%), Mollusca (4%), and Annelida (2%; Fig. 3). For most 18S groups, relative abundances either significantly decreased (*p* < 0.05) with rising temperatures (Dinophyceae, Cryptophyceae, Spirotrichea, and Filosa-Thecofilosea; ρ = –0.26 to –0.83) or were not significantly correlated (*p* > 0.24) with temperature (Mamiellophyceae, Annelida, and Mollusca; Fig. 3). Bacillariophyta abundance increased slightly with temperature (ρ = 0.23, *p* = 0.02), while Syndiniales and Arthropoda exhibited the strongest positive relationships with temperature (ρ = 0.8 and 0.53, *p* < 0.001; Fig. 3). There were 418 18S ASVs (4%) with differentially abundant log2 fold values between cold and warm conditions, with 290 belonging to major groups (Table S2). 18S ASVs exhibited a wide range in log2 fold abundance (–28 to 28) that varied between and within taxonomic groups (Fig. S3). For example, ASVs within Bacillariophyta_X (Bacillariophyta), Gymnodiniales (Dinophyceae), and Tintinnida (Spirotrichea) spanned a wide range of log2 fold values, whereas most ASVs within Syndiniales and Cryptophyceae were expressed in either warm or cold conditions, respectively (Fig. S3).

**Fig. 3:**
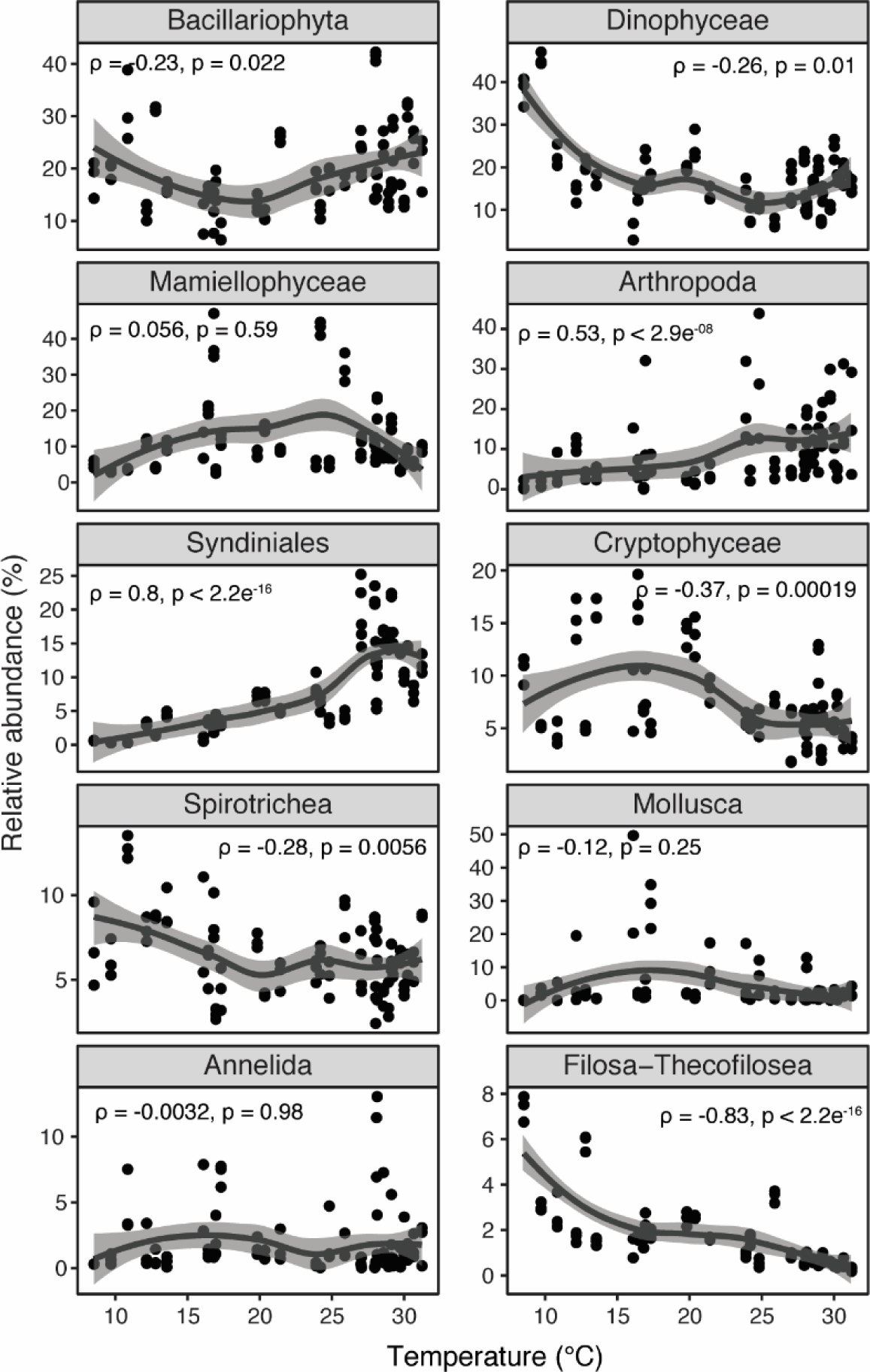
Group-specific temperature effects on 18S relative abundance. 18S relative abundance vs. temperature relationships across all points for major class level groups. Local regression (loess) curves were fit to the data, with solid black lines representing the smoothed trends and shaded gray regions the 95% confidence intervals of the fit. Spearman correlation coefficients (ρ) and *p*-values are indicated in the panels. Abundance values reflect sample triplicates.

### Temperature effects on trophic interactions

We used SpiecEasi to infer microbial interactions (including cross-domain) under warm and cold conditions, combining the most abundant 16S and 18S ASVs (Fig. 4). Networks varied by temperature; the warm network had 991 edges (55% positive) between 360 ASVs, while the cold network revealed 851 edges (59% positive) between 292 ASVs (Table 1). Interactions from both networks were dominated by similar microbes (e.g., Bacillariophyta, Dinophyceae, Alphaproteobacteria, and Gammaproteobacteria), though certain groups, like Syndiniales, contributed more to warm network edges (Fig. 4A, C; Table S3). In the warm network, edges were most common among eukaryotes (Fig. 4A–B), including Bacillariophyta associated with other Bacillariophyta (73 edges), Dinophyceae (62 edges), Spirotrichea (30 edges), and Syndiniales (26 edges), as well as Dinophyceae ASVs associated with Spirotrichea (28 edges), Syndiniales (27 edges), and other Dinophyceae (24 edges). In the cold network, interactions among 18S groups remained common (e.g., Bacillariophyta–Bacillariophyta and Bacillariophyta–Dinophyceae), however, 16S ASVs contributed more to the top interactions (Fig. 4C). Some examples of 16S interactions in the cold network included Alphaproteobacteria associated with Bacteroidia (25 edges), Gammaproteobacteria (21 edges), and Bacillariophyta (16 edges), as well as Gammaproteobacteria associated with Bacillariophyta (21 edges), Bacteroidia (16 edges), and other Gammaproteobacteria (15 edges; Fig. 4D). Though represented by similar class level groups, network properties varied at the domain level (18S vs. 16S) and between warm and cold conditions (Table 1; Table S4). Mean degree (number of edges per ASV) was significantly higher (*p* < 0.001) in the cold vs. warm network for 16S ASVs, while there was no temperature-based difference in mean degree for 18S (Fig. 5A). Within the same network, degree was significantly higher (*p* < 0.0001) for 18S vs. 16S on average (Fig. 5A). Closeness centrality was significantly higher (*p* < 0.001) on average for 18S/16S samples in the cold vs. warm network and for 18S relative to 16S samples in both networks; there was no significant difference (*p* > 0.05) among treatments for betweenness centrality (Table 1).

**Fig. 4:**
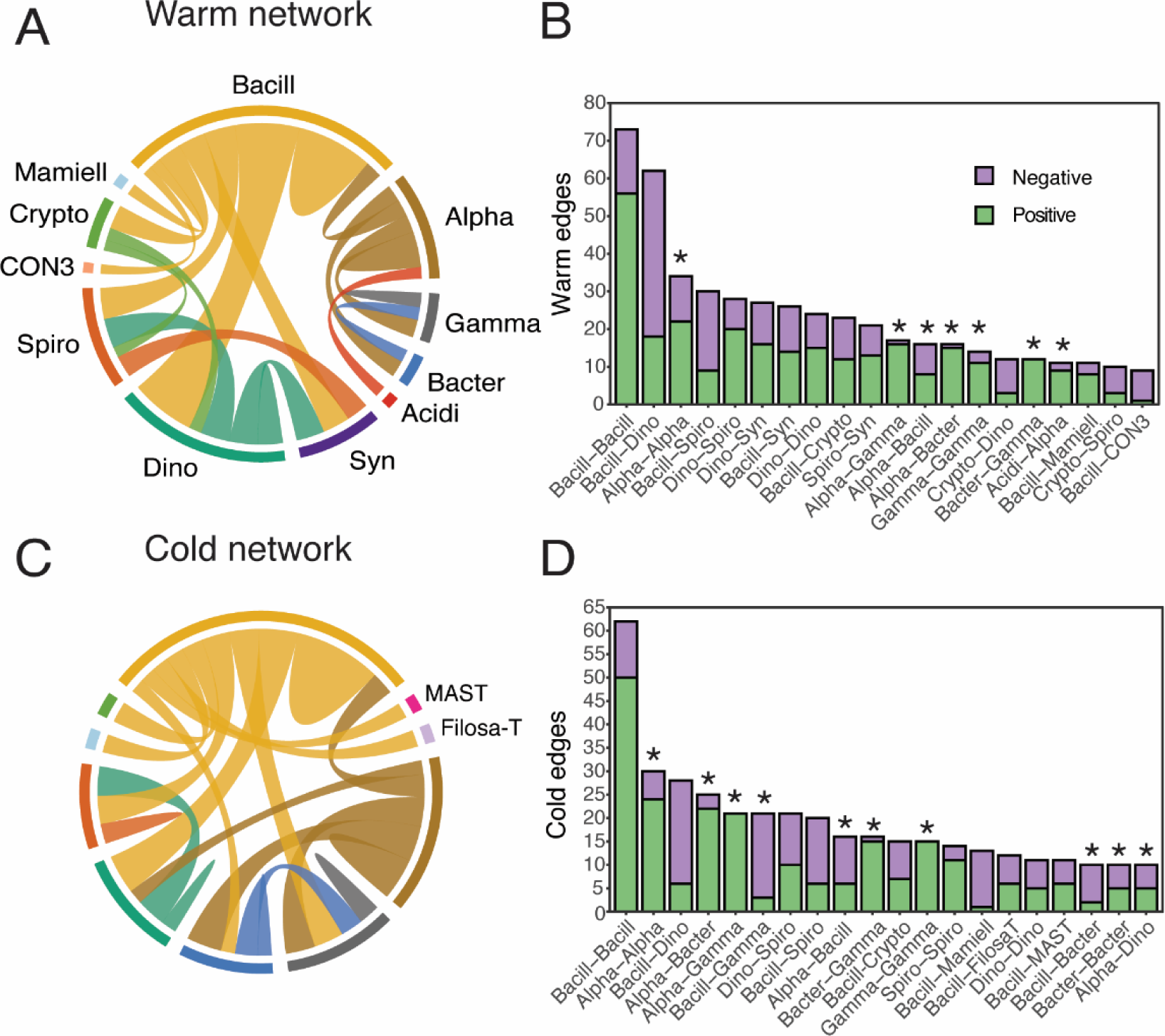
Temperature-based trophic interactions inferred via SpiecEasi. **A, C.** Top 20 most prevalent network associations (edges) between class level 16S and 18S groups (4–60 ASVs per class), representing the major types of interactions in warm (**A**) and cold (**C**) temperatures. Line thickness indicates the number of associations between groups. **B, D**. Total number of positive (green) and negative (light purple) edges for the most abundant pairings (at class level) for warm (**B**) and cold (**D**) networks. Interactions involving 16S groups are distinguished by asterisks. Abbreviations for 18S/16S groups: Bacill = Bacillariophyta; Dino = Dinophyceae; Syn = Syndiniales; Spiro = Spirotrichea; Crypto = Cryptophyceae; Mamiell = Mamiellophyceae; CON3 = CONThreeP; FilosaT = Filosa-Thecofilosea; Alpha = Alphaproteobacteria; Gamma = Gammaproteobacteria; Bacter = Bacteroidia; Acidi = Acidimicrobiia.

**Fig. 5:**
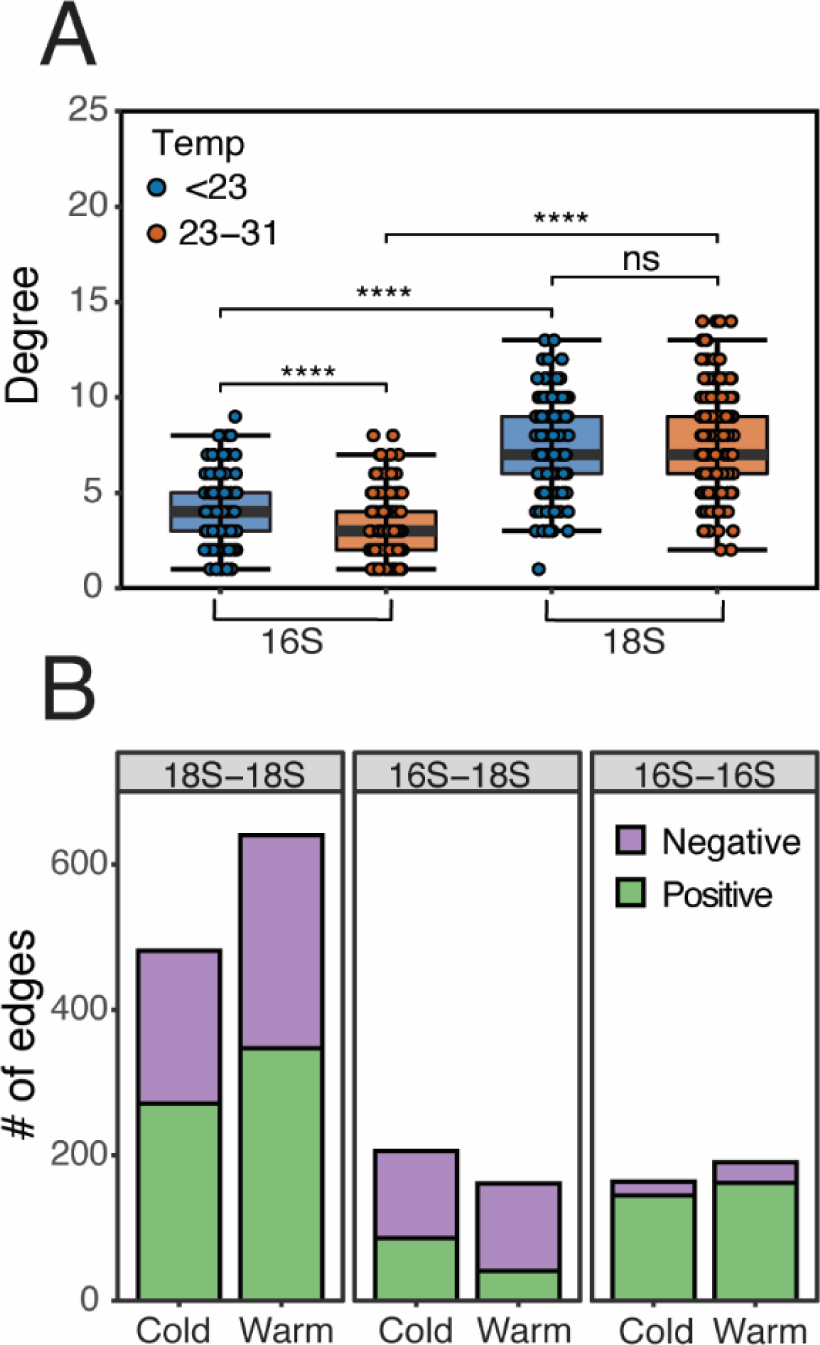
Key properties for warm and cold networks. **A.** Mean degree for all 16S and 18S ASVs in the warm (red) and cold (blue) networks. Degree was compared between temperature networks and domains within the same network (e.g., 18S vs. 16S). Statistical significance is indicated in the figure panels (ns = not significant, **** = *p*-values < 0.0001). **B.** Total number of edges from the warm and cold networks, faceted by kingdom level pairings (18S–18S, 16S–18S, and 16S– 16S). Both positive (green) and negative (light purple) edges are shown for each temperature network and pairing.

**Table 1:** Key properties for cold (<23°C) and warm (23–31°C) networks. Properties include overall number of network edges, as well as mean degree, betweenness centrality, and closeness centrality estimated from all 18S and 16S ASVs in warm and cold networks. Standard deviations are shown in parentheses.

| Property | Warm |  |  | Cold |  |  |
| --- | --- | --- | --- | --- | --- | --- |
|  | 16S | 18S | Total | 16S | 18S | Total |
| Nodes | 170 | 190 | 360 | 133 | 159 | 292 |
| Edges | - | - | 991 | - | - | 851 |
| Degree | 3.18 (1.61) | 7.58 (2.69) | 5.51 (3.14) | 4.02 (1.76) | 7.35 (2.24) | 5.83 (2.63) |
| Betweenness | 0.01 (0.08) | 0.01 (0.01) | 0.01 (0.05) | 0.01 (0.01) | 0.01 (0.01) | 0.01 (0.01) |
| Closeness | 0.24 (0.12) | 0.28 (0.02) | 0.26 (0.08) | 0.26 (0.02) | 0.29 (0.02) | 0.28 (0.03) |

Overall, the number of edges observed among 18S ASVs were elevated (∼33% higher) in warm vs. cold networks, whereas the number of edges observed among 16S ASVs and between domains (16S–18S) were relatively stable (Fig. 5B). A similar proportion (∼55%) of 18S–18S edges were positive in both networks, whereas the proportion of positive interactions increased for 16S–18S (41% vs. 25%) under cold vs. warm conditions (Fig. 5B). Inferred 16S–16S edges were largely positive (∼86%) in both networks (Fig. 5B). Diatom–bacteria interactions accounted for ∼25% of 16S–18S edges in both networks, largely involving chain-forming diatom ASVs (*Thalasiossira*, *Chaetoceros*, *Skeletonema,* and *Rhizosolenia*) associated with ASVs from Flavobacteriales (Bacteroidia) and Rhodobacterales (Alphaproteobacteria). Diatoms interacted more often with members of Gammaproteobacteria (Cellvibrionales and SAR86 Clade) under cold conditions (Table S3). Regardless of temperature, the bulk of 16S–16S interactions involved ASVs from SAR11 Clade associated with Rhodobacterales and Flavobacteriales (Table S3).

## Discussion

Amplicon metabarcoding has expanded our ability to characterize microbial life, with such techniques being non-invasive, holistic (e.g., microbes to metazoans), and sensitive to underrepresented members of the community (Caron and Hu, 2019; Santoferrara *et al.*, 2020). The inclusion of metabarcoding with temporal (or spatial) sampling has established reliable drivers of microbial richness, relative abundance, and composition, chief among them surface water temperature (Fuhrman *et al.*, 2015; Sunagawa *et al.*, 2015; Ibarbalz *et al.*, 2019). Given the expected rise of ocean temperatures, it remains critical to better understand the consequences of warming on marine microbes and their interactions with other organisms (Caron and Hutchins, 2013; Hutchins and Fu, 2017), which have so far been challenging to predict.

Here, we employed 16S and 18S rRNA metabarcoding from samples collected over a full year in the Skidaway River Estuary. To alleviate issues in interpreting metabarcoding data, including gene copy number (Caron *et al.*, 2012), we transformed to relative abundance, processed 16S and 18S sets separately, and assessed temperature-based responses within a given taxonomic tank. Over a wide temperature gradient (9–31°C), we observed shifts in microbial (and metazoan) population dynamics and temperature-based interactions. Baseline information on the response of microbes to rising temperature is important (Cavicchioli *et al.*, 2019), as even slight changes in temperature can influence microbial communities and associated ecosystem-level processes.

Over the course of our yearly sampling, ∼75% of total 16S relative abundance was attributed to Alphaproteobacteria, Gammaproteobacteria, and Bacteroidia, all of which are among the most abundant bacteria groups in coastal environments (Buchan *et al.*, 2014). Beyond an increase in production and respiration rates (Sarmento *et al.*, 2010), some studies suggest warming may favor copiotrophic bacteria (e.g., Rhodobacterales or Flavobacteriales) that can better exploit temperature-based changes in phytoplankton-derived organic carbon (von Scheibner *et al.*, 2014; Arandia-Gorostidi *et al.*, 2017b). We found that Alphaproteobacteria, largely represented by SAR11 Clade and Rhodobacterales, decreased in relative abundance with rising temperatures. Though present globally, SAR11 may be more susceptible to warming, related to their streamlined genomes and poor regulation of certain metabolic genes (Giovannoni *et al.*, 2014; Giovannoni, 2017). Other major 16S groups, like Bacteroidia (mainly Flavobacteriales) and Gammaproteobacteria (e.g., SAR86 Clade and Cellvibrionales) peaked in relative abundance at 18–25°C, decreasing thereafter. Though within the context of spatial sampling, similar trends have been observed for bulk 16S richness, with highest values occurring at intermediate temperatures (∼15°C) and steadily decreasing beyond that (Sunagawa *et al.*, 2015).

Albeit less abundant in our survey, we identified significant temperature-abundance relationships among other ecologically important bacterial groups. For instance, relative abundance of Acidimicrobiia (Actinomarinales and Microtrichales) and Deltaproteobacteria (e.g., Bdellovibrionales and Desulfobacterales) exhibited strong and linear increases with rising temperatures, while relative abundance increased exponentially for the archaea group, Nitrososphaeria (Nitrosopumiales). Anaerobic microbes like Desulfobacterales (sulfate-reducing) and Nitrosopumiales (ammonia-oxidizing) occupy important roles in sulfur and nitrogen cycling within coastal estuaries and have been found (via gene libraries and transcriptomics) to be prevalent in marine waters and sediments off Georgia (Lasher *et al.*, 2009; Hollibaugh *et al.*, 2011, 2014). Oxyphotobacteria (notably Synechococcales) also increased exponentially, consistent with some model predictions that favor growth of smaller-sized cyanobacteria under higher temperatures (Feng *et al.*, 2009). Temperature-abundance relationships may be more pronounced among less abundant groups due to lower functional diversity compared to others (e.g., Alpha- or Gammaproteobacteria). Still, it is important to consider temperature effects across different 16S taxonomic groups, even those less abundant, as these groups encompass diverse metabolic strategies (e.g., heterotrophy, autotrophy, sulfate-reducing, and ammonia-oxidizing) that influence biogeochemical cycling in estuarine food webs.

Eukaryotes in the estuary were dominated (in terms of relative abundance) by Bacillariophyta, Mamiellophyceae, and Dinophyceae, consistent with other coastal 18S barcoding studies (Massana *et al.*, 2015; Tragin *et al.*, 2018; Gong *et al.*, 2020). These protist groups (and many others) are functionally and phylogenetically diverse (Caron *et al.*, 2012; Tragin and Vaulot, 2019), which may explain why we observed little fluctuation in class level relative abundance with temperature. Diatoms are known to exhibit species-specific temperature–growth relationships and occupy thermally driven environmental niches (Canesi and Rynearson, 2016; Rynearson *et al.*, 2020). Moreover, phytoplankton blooms (diatoms and dinoflagellates) have been observed throughout most of the year in the estuary (Verity and Borkman, 2010), indicating potential acclimation of resident phytoplankton communities to varying temperatures. It may also be true that in highly dynamic estuaries, like Skidaway River, other abiotic factors (e.g., tidal flow and vertical mixing) and those which covary with temperature (e.g., nutrients) are important in structuring phytoplankton blooms (Anderson *et al.*, 2018), even more so than temperature at times (Busseni *et al.*, 2020), and should be considered as microbial drivers.

Among major 18S groups, only relative abundances of Syndiniales and Arthropoda had strong and positive relationships with temperature. Both taxonomic groups are ubiquitous in marine systems and contribute to carbon cycling in different ways. Syndinales are obligate parasites that infect a range of hosts (e.g., dinoflagellates, ciliates, and metazoans), often leading to bloom termination and the release of dissolved organic carbon (Guillou *et al.*, 2008; Velo-Suárez *et al.*, 2013). Arthropoda, notably copepods (*Acartia* and *Euterpina*) at our site, represent important trophic links in estuarine systems (Tan *et al.*, 2004), consuming phytoplankton (and microzooplankton) biomass that can be transferred to higher trophic levels (Steinberg and Landry, 2017). Increased relative abundance of Syndiniales and Arthropods in our system may reflect underlying metabolic processes that are more favorable (e.g., heterotrophy vs. autotrophy) under warmer conditions (Chen *et al.*, 2012). While temperature may influence parasitic infection and prey ingestion, these processes also depend on host/prey qualities (e.g., density, composition, or nutritional state), making it difficult to parse direct temperature effects (Lawrence and Menden-Deuer, 2012). Other heterotrophic protist groups in our survey (e.g., Spirotrichea and Filosa-Thecofilosea) decreased in relative abundance with temperature, owing to the complex nature in predicting temperature response based purely on functional strategies.

Across both 16S and 18S samples, <5% of ASVs were differentially abundant (via log2 fold abundance) in cold vs. warm conditions. A similar finding was recently observed for coastal microbes incubated at temperatures ∼3°C above ambient (Wang *et al.*, 2021). This suggests that at the ASV level, most microbes (especially bacteria which did not cluster by temperature) were distributed across a range of temperatures and adapted to environmental variability in this system. Our findings represent only a single year; however, microbial dynamics are often repeatable (Ward *et al.*, 2017), including in the Skidaway River, where yearly temperatures (6– 33°C) are consistent over interannual scales (Verity, 2002). Temperature effects we observed may also be related to microbial biogeography, as inshore communities are thought to be less susceptible to temperature variability compared to those offshore, where conditions are more stable (Wang *et al.*, 2021); though extreme warming beyond yearly temperature maxima may elicit shifts in coastal microbes (see Kling *et al.*, 2020).

Marine microbial communities are governed by complex trophic interactions (e.g., predation, competition, and mutualism), influencing food web dynamics and the fate of organic carbon (Fuhrman *et al.*, 2015; Worden *et al.*, 2015). To investigate microbial interactions, we used correlation network analysis, a technique that has become increasingly applied to metabarcoding data to infer significant associations between co-occurring ASVs (Faust and Raes, 2012).

Interactions inferred via sequence networks represent a baseline of information (Santoferrara *et al.*, 2020), spurring future work (and verification) from quantitative methods like microscopy or cell sorting. Prior temporal networks have included biotic and abiotic features (including temperature) to distinguish species-environment dynamics (Cram *et al.*, 2015; Needham and Fuhrman, 2016); however in this study, we focused on assessing only biological interactions with respect to distinct temperature conditions (<23 vs. 23–31°C). At a broad level, warm and cold networks appeared similar, dominated by the most relatively abundant 16S and 18S groups (e.g., Bacillariophyta, Dinophyceae, Alpha-, and Gammaproteobacteria). However, we observed temperature-specific differences in network properties (e.g., degree and centrality) and interactions between certain taxonomic groups, which may have implications for how we predict future temperature effects on marine microbes and their trophic interactions.

We observed considerable variation in 18S–18S interactions between temperatures, becoming more abundant in warmer conditions. This increase was mainly attributed to Bacillariophyta– Dinophyceae interactions, involving unassigned Peridiniales and Gymnodiniales associated with chain-forming diatoms (e.g., *Chaetoceros* and *Thalassiosira*). Heterotrophic dinoflagellates are known to ingest large diatoms (Sherr and Sherr, 2007), though our interactions were often negative (>70%), indicating mutual exclusion (i.e., likely not predation). Negative relationships we observed may reflect phytoplankton mechanical or chemical defenses against predation, especially as larger chain-forming diatoms may be less palatable to smaller grazers (reviewed in Pančić and Kiørboe, 2018). Other groups that became more prominent under warmer temperatures included Syndiniales parasites, having associations with known (dinoflagellates and ciliates) and putative (diatoms) hosts (Sassenhagen *et al.*, 2020; Vincent and Bowler, 2020). We recognize the higher number of warm interactions among eukaryotes may be driven by a slight increase in 18S ASVs (or nodes) included in the warm (190) relative to the cold network (159). Even so, network properties for 18S groups (e.g., average degree) did not vary by temperature, suggesting their contribution to network connectivity was conserved across temperature regimes in this system.

Though contributing fewer edges compared to eukaryotes, 16S ASVs became more prominent among the top interactions in the cold network and contributed more to network connectivity (higher degree and centrality). Diatom–bacteria interactions accounted for ∼25% of cross domain (16S–18S) relationships (in both networks), consistent with the well-established importance of these interactions in aquatic systems (Amin *et al.*, 2012). Diatoms in our study were most often associated (both positively and negatively) with Rhodobacterales and Flavobacteriales, opportunistic bacteria known to rapidly exploit diatom-produced organic matter (Teeling *et al.*, 2012; Buchan *et al.*, 2014). Heterotrophic bacteria depend on photosynthetic production to sustain population growth, while in turn, phytoplankton rely on bacteria-derived metabolites and other compounds (e.g., nutrients, vitamins, and hormones) that are exchanged at the cellular level (Amin *et al.*, 2012, 2015; Moran *et al.*, 2012; Seymour *et al.*, 2017). Mutualistic diatom–bacteria interactions may be favored under warmer temperatures (Arandia-Gorostidi *et al.*, 2017a), though we did not observe evidence of this in our temperature-based networks. Interestingly, interactions between diatoms and members of Gammaproteobacteria (SAR86 Clade and Cellvibrionales) increased in cold temperatures, with such interactions being mostly negative (90%) and indicative of potentially antagonistic or competitive relationships (e.g., competition for dissolved organic matter) or niche partitioning. In general, higher connectivity of 16S ASVs in cold conditions may reflect strategies among resident bacteria to interact with diatoms (or other microbes), exchanging (or competing for) metabolites needed for growth.

Compared to cross-domain interactions, which included both positive and negative edges, 16S– 16S interactions were overwhelmingly positive in both networks (86%), indicating possible mutualism among bacteria groups (though environmental niches may also overlap). The most dominant 16S–16S interactions involved SAR11 and either Rhodobacterales or Flavobacteriales.

Oligotrophic (and genetically streamlined) microorganisms like SAR11 may rely on copiotrophic bacteria (or phytoplankton) to assimilate sources of carbon or other metabolites necessary for growth (Buchan *et al.*, 2014; Giovannoni *et al.*, 2014), resulting in the expansion of their environmental niches in dynamic coastal habitats. These types of bacterial interactions, including those between SAR11 and Rhodobacterales, are often saturated in free-living (0.22–3 µm) communities and may be underestimated here in our particle-attached (<200 µm) samples (Milici *et al.*, 2016). In relation to temperature, Arandia-Gorostidi *et al.* (2017a) found a larger increase in bacterial abundance and bacteria–phytoplankton carbon transfer in the particle-attached vs. free living fraction under warming conditions. Nevertheless, there is merit to considering multiple size fractions within microbial communities, in this case offering deeper insight into the response of microbes to temperature (or other environmental) variability.

In summary, we applied temporal metabarcoding of multiple gene regions (18S and 16S rRNA) in a subtropical estuary, revealing group-specific trends in relative abundance with respect to temperature. Certain microbes (e.g., Deltaproteobacteria, Acidimicrobiia, and Syndiniales) increased in relative abundance with rising temperatures, whereas other relatively abundant groups (e.g., Alpha- and Gammaproteobacteria, Bacillariophyta, and Dinophyceae) decreased, became saturated, or were stable. We constructed temperature-specific covariance networks, revealing increased 18S–18S interactions under warm temperatures (e.g., Bacillariophyta– Dinophyceae) and the emerging contribution of 16S ASVs to interactions and network connectivity under cold conditions. Future temperature-based work will benefit from additional quantitative (e.g., qPCR or cell counting) approaches to verify trends in microbial population dynamics observed here (Santoferrara *et al.*, 2020). Analysis of rRNA/rDNA ratios or gene expression patterns (Alexander *et al.*, 2015; Hu *et al.*, 2018) will also be important in the context of temperature variability, providing information on community members or genes that may be up- or down-regulated in different temperatures. There is also a need to promote higher sampling resolution and extended time series work (over multiple years) in coastal and open ocean regions (from tropical to polar), improving on the robustness of temperature effects observed here and strengthening model predictions on biogeochemistry. Monitoring how microbial communities respond to current temperature fluctuations will help us to predict their future responses, particularly important given the role of microbes (and their interactions) in driving ecosystem productivity, trophic transfer, and carbon cycling.

## Experimental procedures

### Sample collection

Surface layer (1 m) water samples were collected on a weekly to monthly basis over a full year in the Skidaway River Estuary (latitude, 31°59′25.7″N; longitude, 81°01′19.7″W), encompassing 33 sampling days. Sampling always occurred at high tide to normalize effects of semidiurnal tides. Samples were collected with Niskin bottles, filtered through 200-μm mesh, and transferred to a nearby lab for processing. Water samples (250–1000 ml) were filtered in triplicate through 47-mm, 0.22-μm polycarbonate filters (Millipore) using gentle vacuum filtration and stored at – 80°C. The DNeasy PowerSoil kit (Qiagen) was used to extract total DNA, including whole microbial cells and environmental DNA that passed through the 200-μm filter, following manufacturer’s protocols. DNA samples were eluted in 10 mM Tris–HCl (pH = 8.5). DNA concentrations were assessed using a Qubit dsDNA HS kit (Thermo Scientific), ranging from 2– 5 ng μl^-1^ per sample. Environmental factors (e.g., temperature, salinity, nutrients, and chlorophyll) were measured concurrently with metabarcoding samples (presented in Anderson and Harvey 2020). Here, we used temperature data, measured with a YSI probe (YSI 600 QS) and included other important environmental data (e.g., total dissolved nutrients and chlorophyll) to provide context for the sampling site (Fig. 1A).

### PCR conditions and DNA sequencing

We employed a two-step PCR approach with two different primer sets, targeting bacteria and archaea (16S rRNA), as well as protists and some metazoans (18S rRNA). We used primers from Stoeck *et al.* (2010), targeting the V4 region of the 18S rRNA gene: forward (5′-CCAGCASCYGCGGTAATTCC-3′) and reverse (5′-ACTTTCGTTCTTGATYRA-3′). Primers from Parada *et al.* (2016) were used to target the V4–V5 region of the 16S rRNA gene: forward (5′-GTGYCAGCMGCCGCGGTAA-3’) and reverse (5′-CCGYCAATTYMTTTRAGTTT-3’). Sequencing of the 16S V4–V5 region has been shown to better represent marine microbial diversity and resolve taxa that may be underestimated (e.g., SAR11) by shorter amplicon reads (Parada *et al.*, 2016). PCR conditions for 18S involved an initial denaturation step at 98°C for 2 min, 10 cycles of 98°C for 10 s, 53°C for 30 s, and 72°C for 30 s, followed by 15 cycles of 98°C for 10 s, 48°C for 30 s, and 72°C for 30 s, and a final extension of 72°C for 2 min (Stoeck *et al.*, 2010; Hu *et al.*, 2015). For 16S, PCR conditions consisted of an initial denaturation of 95°C for 2 min, 25 cycles of 95°C for 45 s, 50°C for 45 s, and 68°C for 90 s, followed by a final elongation step of 68°C for 5 min (Parada *et al.*, 2016). PCR products from the initial 16S or 18S run were purified and size-selected using AMPure XP Beads (Beckman Coulter, A63881). A second PCR step was carried out with dual Illumina indices. Two separate amplicon sequencing runs (16S and 18S) were performed using an Illumina MiSeq (2 × 250 bp) at the Georgia Genomics and Bioinformatics Core at the University of Georgia.

### Bioinformatics analysis

Demultiplexed 16S and 18S FASTQ files were imported and processed separately in QIIME 2 (Bolyen *et al.*, 2019). Amplicon sequence variants (ASVs) were inferred using paired-end DADA2 (Callahan *et al.*, 2016). Truncation lengths of forward and reverse reads and primer trimming (e.g., trim-left or trim-right) were defined based on read-quality profiles. Default DADA2 parameters (e.g., maxEE) were used for other flags. Taxonomic annotation of 18S ASVs was performed using QIIME 2 compatible files from the Protist Ribosomal Reference (PR2) database (Version 4.12.0; Guillou *et al.*, 2013), while 16S taxonomy was assigned using the SILVA database (Version 138; Pruesse *et al.*, 2007). For both gene regions, a Naïve Bayes Classifier was used to train the sequences against the reference databases using the feature-classifier plugin in QIIME 2 (Bokulich *et al.*, 2018). QIIME 2 taxonomy and count table artifact files (.qza files) were imported into R (Version 3.6.3; https://cran.r-project.org) using the read_qza function from the qiime2R package (https://github.com/jbisanz/qiime2R).

Sequences were deposited at the Sequence Read Archive of the National Center for Biotechnology Information (NCBI) and made publicly available under accession numbers PRJNA575563 (18S) and PRJNA680039 (16S). R code used for downstream ASV analyses and visualizations are on GitHub (https://github.com/sra34/SkIO-temp), including QIIME 2 output files (count, taxonomy, and representative sequences), metadata tables, and network-related files.

### Statistical analyses

We used a variety of R packages, including phyloseq (McMurdie and Holmes, 2013), vegan (Oksanen *et al.*, 2018), and DESeq2 (Love *et al.*, 2014) to explore temperature effects on 16S and 18S communities. We focused on comparing microbial community dynamics between two distinct temperature conditions, considered here as cold (<23°C; 14 sampling points) or warm (23–31°C; 19 sampling points). 18S ASVs that were unassigned at the supergroup level were removed, while 16S ASVs that were unassigned at the kingdom level or assigned to either Chloroplast (order level) or Mitochondria (family level) were removed before further processing. After these filtering steps, we ended up with 5,976,579 total sequences for 18S (28,912– 104,151), assigned to 9,702 ASVs. There were 7,629,573 total sequences for 16S (47,250– 177,521), assigned to 15,716 ASVs. Rarefaction curves were observed for both 16S and 18S with the ggrare function in the R package ranacapa (https://github.com/gauravsk/ranacapa). Taxonomic assignments and unfiltered read counts for 18S and 16S ASVs over the sampling period are included in Table S1.

We rarefied samples to the minimum read count for 16S (47,250 reads) and 18S (28,912 reads) to account for variability in the number of reads per sample. Alpha diversity metrics (observed richness and Shannon diversity) were estimated with the estimate_richness function in phyloseq. Shapiro–Wilks normality tests were performed on the diversity data, revealing a parametric distribution; *t*-tests were used to test significant differences in diversity between cold and warm conditions (adjusted Holm *p*-values < 0.05). Non-metric dimensional scaling (NMDS) was performed using Bray–Curtis distances, following the removal of global singletons (ASVs found once across 33 d) and transformation of sequence reads to relative abundance. Analysis of similarity (ANOSIM) tests were used to verify significant differences in composition between temperature conditions. We used the DESeq function in DESeq2 to identify 16S or 18S ASVs that were differentially abundant between cold and warm conditions. Logarithmic fold changes (log2 fold) were determined and filtered for statistical significance (adjusted *p*-values < 0.001). Relative abundance vs. temperature relationships for 16S and 18S groups at the class level were assessed with local regression (loess) curves, distinguishing smoothed trends (both linear and non-linear) in relative abundance over a wide temperature gradient. Spearman correlations (ρ) were estimated from these relationships using the ggpubr package in R (https://github.com/kassambara/ggpubr). Group-specific temperature analyses focused on 16S or 18S taxa that were most abundant in each dataset (at class level), defined here as groups with an average relative abundance ≥1% over all samples.

### Covariance networks

Temperature-based trophic interactions were inferred using the SParse InversE Covariance estimation for Ecological ASsociation Inference (SpiecEasi; Version 1.1.0) package in R (Kurtz *et al.*, 2015), which allows for cross-domain interactions to be determined from a merged ASV count table (Tipton *et al.*, 2018). Unlike other modes of network inference that infer pairwise correlations (e.g., Spearman, SparCC, and CoNet), SpiecEasi infers direct associations using conditional independence and computes an inverse covariance matrix (Kurtz *et al.*, 2015; Röttjers and Faust, 2018). SpiecEasi is robust to the compositional nature of amplicon sequencing data, resulting in sparser networks that limit spurious ASV correlations (Kurtz *et al.*, 2015).

In preparation for network analysis, 16S and 18S samples were separated into phyloseq objects based on warm vs. cold temperatures. To reduce ambiguous relationships and improve sparsity, we removed ASVs that were unassigned at the class level and trimmed each dataset to include only ASVs with >5 read counts in at least 50% or 10% of samples for 18S and 16S, respectively. Networks were constructed with the spiec.easi function using 16S and 18S ASV count tables as input (with matching sample IDs) and the “mb” (Meinshausen–Buhlmann) neighborhood selection setting. The Stability Approach to Regularization Selection (StARS) was used to select the optimal sparsity parameter with a threshold set to 0.05 (Liu *et al.*, 2010). The spiec.easi function performs center log normalization of ASV count tables, avoiding pre-transformation of data (Tipton *et al.*, 2018). The output of SpiecEasi is a correlation matrix of positive and negative values (weights) for all significantly paired edges.

Networks were visualized in Cytoscape (Shannon *et al.*, 2003) and summary statistics such as degree, betweenness centrality, and closeness centrality were estimated with the NetworkAnalyzer plugin. Network degree refers to the number of edges connected to a given node (or ASV), whereas betweenness and closeness centrality indicate either the sum of the shortest paths that move through an ASV or the proximity of a given ASV to all other ASVs in the network; higher centrality indicates a greater contribution to network connectivity (Röttjers and Faust, 2018). Differences in network topology (mean degree and centrality) were assessed at the domain level (16S vs. 18S) and between temperatures (e.g., 16S warm vs. 16S cold) using pairwise Wilcoxon tests (*p* < 0.001). To characterize the most important interactions in each network, we filtered networks to include the top 20 most prevalent class level interactions (16S and/or 18S), distinguishing between the number of positive and negative edges. Major network associations were visualized using chord diagrams in the R package circlize (Gu *et al.*, 2014).

## Acknowledgements

The authors would like to thank Tina Walters (UGA/SkIO) for help with initial PCR processing of metabarcoding samples. We thank Luke Thompson (NOAA/NGI) for helpful review of the manuscript. This work is funded by a Sloan Research Fellowship to E.L.H., and a National Science Foundation Grant (OCE-1831625) that supported the collaboration of S.R.A., M.C., and E.L.H. during the summer of 2020.

## Supporting information

**Table S1:** Taxonomic assignments (kingdom to species) and raw sequence counts per sampling day for all assigned 16S and 18S ASVs (separate sheets). Read counts have not been rarefied. 16S ASVs are labeled as “bASV” to denote bacteria from eukaryotes.

**Table S2:** Log2 fold change values based on significantly differential (*p* < 0.001) read abundance for 16S and 18S ASVs (separate sheets) between cold and warm samples. Taxonomic annotation (kingdom-through species), *p*-value, and adjusted *p*-value (p-adj) are shown for each differentially abundant ASV.

**Table S3:** Information for all ASV-ASV edges (16S and 18S) inferred via the warm and cold network (separate sheets). Edge information includes taxonomic annotation conserved across loci (kingdom, class-through species) per node per pairing (Node 1 and Node 2), edge weight, sign (negative or positive), and interaction type (e.g., 16S–18S or 18S–18S). All edges were significant.

**Table S4:** Network properties for each 16S and 18S ASV (kingdom, class-through species annotation) present in the warm or cold network (separate sheets). Properties include degree, closeness centrality, and betweenness centrality.

**Fig. S1:** Rarefaction curves of species richness vs. sequence read abundance across all sampling days for 16S (left) and 18S (right). Curves were estimated using a step size of 100.

**Fig. S2:** Log2 fold change of 16S ASVs with significantly differential abundance (adjusted *p*-values < 0.001) under cold vs. warm conditions. Each point represents an individual ASV and ASVs are grouped by order (left y-axis) and class (right y-axis) level taxonomy.

**Fig. S3:** Log2 fold change of 18S ASVs with significantly differential abundance (adjusted *p*-values < 0.001) under cold vs. warm conditions. Each point represents an individual ASV and ASVs are grouped by order (left y-axis) and class (right y-axis) level taxonomy.

