## Supplemental Figures for "Investigating temperature effects on coastal microbial populations and trophic interactions with 16S and 18S rRNA metabarcoding"

Running title: Temperature-based microbial interactions.

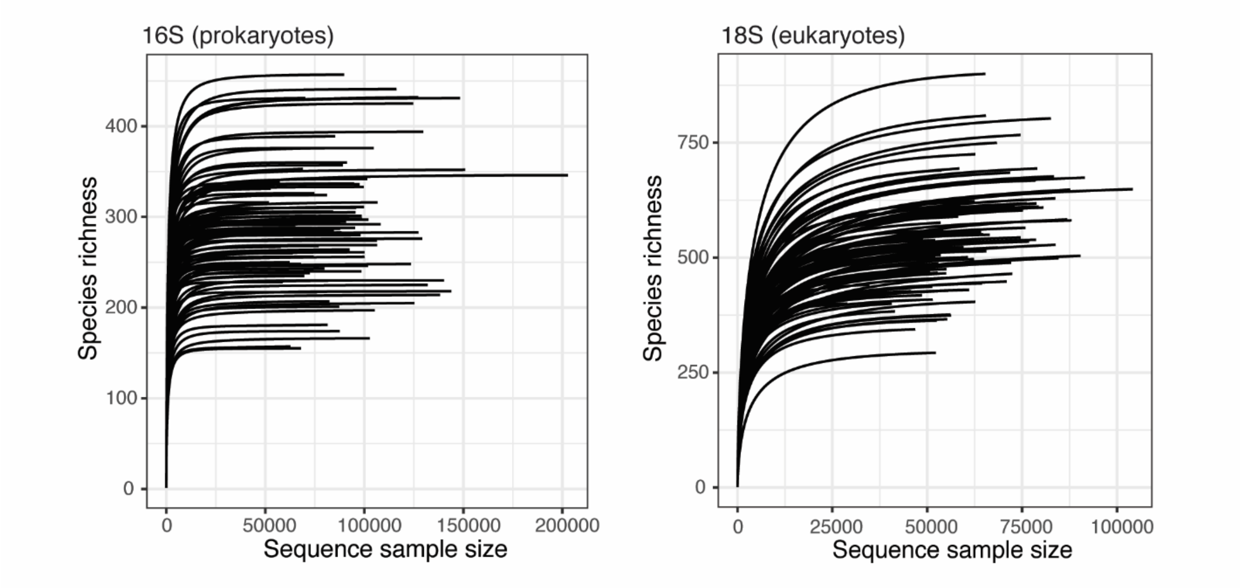


**Fig. S1**: Rarefaction curves of species richness vs. sequence read abundance across all sampling days for 16S (left) and 18S (right). Curves were estimated using a step size of 100.

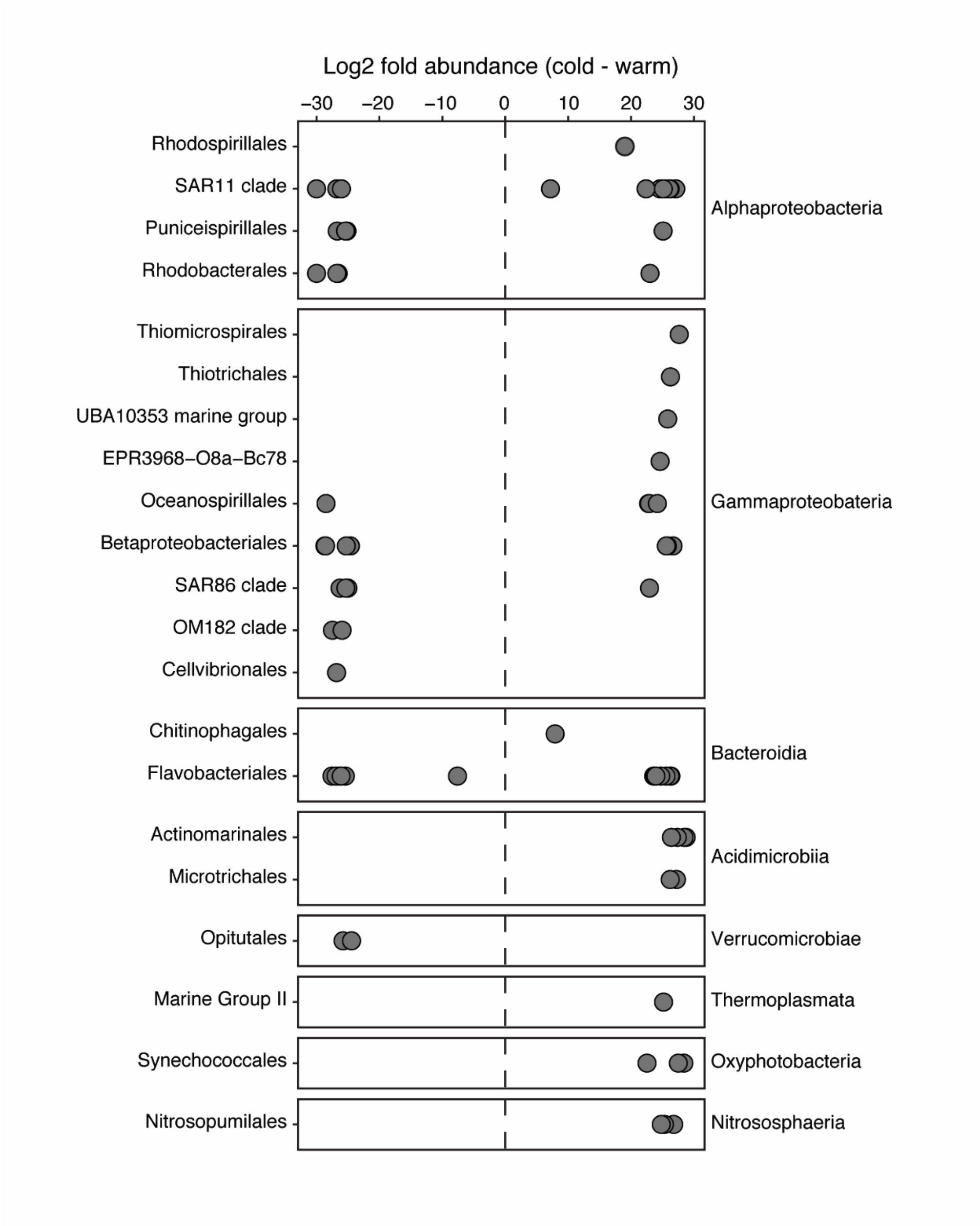


**Fig. S2**: Log2 fold change of 16S ASVs with significantly differential abundance (adjusted *p*-values < 0.001) under cold vs. warm conditions. Each point represents an individual ASV and ASVs are grouped by order (left y-axis) and class (right y-axis) level taxonomy.

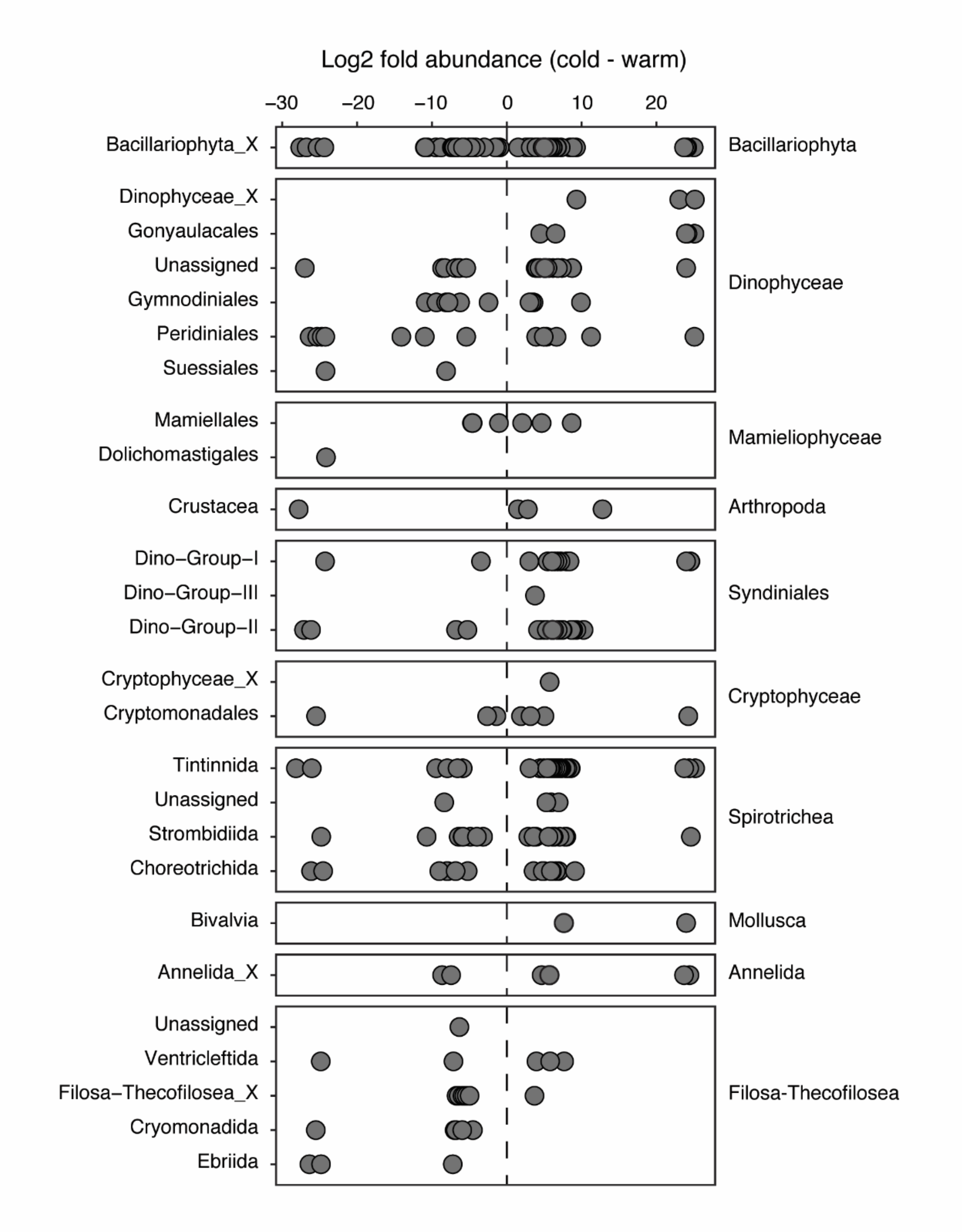


**Fig. S3**: Log2 fold change of 18S ASVs with significantly differential abundance (adjusted *p*-values < 0.001) under cold vs. warm conditions. Each point represents an individual ASV and ASVs are grouped by order (left y-axis) and class (right y-axis) level taxonomy.
